# Efficient simultaneous mutagenesis of multiple genes in specific plant tissues by multiplex CRISPR

**DOI:** 10.1101/2020.11.13.381046

**Authors:** Norbert Bollier, Rafael Andrade Buono, Thomas B. Jacobs, Moritz K. Nowack

## Abstract

Multiplex CRISPR approaches enable mutating multiple genes in plants, however it is unclear how feasible this is in tissue-specific mutagenesis. Here we simultaneously mutated six genes either ubiquitously or exclusively in the root cap of Arabidopsis. The mutation frequencies for all target genes were positively correlated and unaffected by the order of gRNAs in the vector, indicating that efficient higher-order mutagenesis in specific plant tissues can be readily achieved.

## Main

Clustered regularly interspaced short palindromic repeats (CRISPR) technology is an established tool for the generation of knockout plants (Zhang *et al*., 2019). Mutagenesis with the CRISPR-associated 9 (Cas9) protein is routinely used to generate stable and inheritable mutant alleles for reverse genetics approaches (Mao *et al*., 2019). Despite its success, limitations remain. First, the manipulation of individual genes may fail to produce phenotypes for groups of genes with redundant or synergistic functions. While this has been partially addressed by multiplexing guide RNAs (gRNAs), there is the concern that as the number of gene targets increases, the chances of obtaining efficiently mutated individuals diminish (Zhang *et al*., 2016). Second, knocking out fundamentally important genes can lead to severe pleiotropic phenotypes or lethality. Tissue-specific knockout of genes in somatic tissues can overcome this limitation (Decaestecker *et al*., 2019; Wang *et al*., 2020; Liang *et al*., 2019). However, the efficiency of simultaneously targeting more than three genes in a tissue-specific context has remained unexplored. Here, by multiplexing gRNAs in *Arabidopsis thaliana* plants expressing Cas9 either ubiquitously *(pPcUBI)* or root cap-specifically *(pSMB)*, we show that six genes can be simultaneously mutated with high efficiency, generating high-order mutant phenotypes already in the first transgenic generation (T1). The mutation frequencies for all target genes were positively correlated and unaffected by the order of the gRNAs in the vector, showing that efficient higher-order mutagenesis in specific plant tissues can be readily achieved.

We selected six efficient gRNAs (Decaestecker *et al*., 2019 and unpublished results) to target the coding sequences of six genes (the reporter gene *GFP*, and the Arabidopsis genes *SMB, EXI1, GL1, ARF7*, and *ARF19*) (Figure 1a) whose loss-of-function lead to easy-to-score phenotypes in T1 seedlings *(gfp*: loss of GFP signal, *smb*: accumulation of living or dead root cap cells (Fendrych *et al*., 2014), *gl1*: absence of trichomes in leaves (Herman *et al*., 1989)) and do not severely affect plant growth or reproduction. Since it has been suggested that there may be a position effect within a gRNA array (Kurata *et al*., 2018) we generated two vectors (hereafter, *pPcUBI(I)* and *pPcUBI(II))* combining Cas9-mTagBFP2 driven by the ubiquitous *pPcUBI* promoter and the six gRNAs in an inverted order (Figure 1b) and transformed these into an Arabidopsis line with ubiquitous expression of a nuclear-localized GFP *(pHTR5:NLS-GFP-GUS* (Decaestecker *et al*., 2019) hereafter, NLS-GFP).

**Figure 1:**
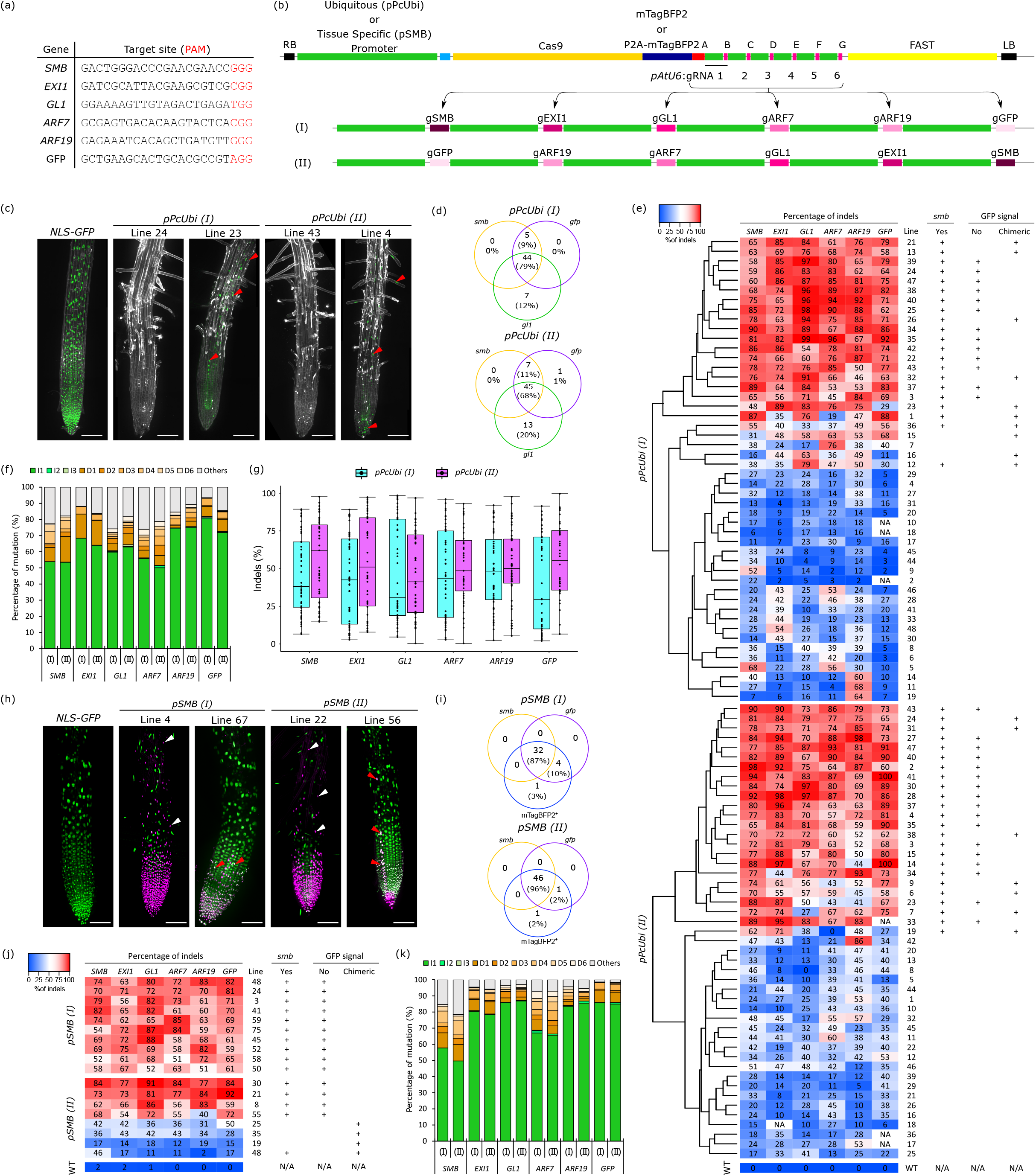
Ubiquitous and root-cap-specific knockout of 6 genes in T1 via CRISPR and CRISPR-TSKO. **a**, gRNA target sequences. PAM sequences are in red. **b**, Schematic diagram of the CRISPR *(pPcUBI)* and CRISPR-TSKO (pSMB) vectors. In addition to their specific promoter, both vectors contain a cloning linker, a G7 terminator and six AtU6-26 promoter driving the expression of each gRNA. The *pPcUBI* vector contain the *SpCas9* nuclease in frame with an *mTagBFP2* fluorescent reporter and the *pSMB* vector contain the *SpCas9* fused to the *mTagBFP2-NLS* coding sequence via a P2A ribosomal skipping peptide. FAST indicates the FAST screenable marker cassette, LB and RB are the left and right borders of the T-DNA region. The six gRNAs were cloned in an inverted order, (I) and (II) **c**, Maximum intensity projections of root tips of a representative NLS-GFP seedling, two *pPcUBI(I)* and two *pPcUBI(II)* T1 seedlings showing the complete (left) and chimeric (right) absence of GFP signal and *smb* mutant phenotype with remnants of root cap cell corpses attached to the root surface. GFP is shown in green and propidium iodide (PI) staining in grey. Red arrowheads indicates a patch of root cells still expressing GFP. Scale bars, 100 μm. **d**, Venn diagram showing the number of plants displaying *smb, gfp*, and *gl1* mutant phenotype in 96 *pPcUBI(I)* (top) and 95 *pPcUBI(II)* (bottom) T1 seedlings. **e**, Genotype analysis by amplicon sequencing. The Heat map shows the percentage of indels for each target site. The hierarchical clustering was done using Heatmapper (Babicki et al. 2016). The phenotype observed for the corresponding seedling is indicated on the right panel. **f**, Frequency of the main mutation types in both *pPcUBI(I)* and *pPcUBI(II)* plants, for the six target sites. I1 to I3: 1- to 3-bp insertion, D1 to D6: 1- to 6-bp deletion, Others: bigger deletions (>6-bp), insertions (> 3-bp) or more complex repair outcomes containing both insertions and deletions. **g**, Percentage of indels observed for the 6 genes targeted in *pPcUBI(I)* and *pPcUBI(II)* T1 plants. **h**, Maximum intensity projections of root tips of a representative NLS-GFP seedling, two *pSMB(I)* and two *pSMB(II)* T1 seedlings grown on 1μM brassinazole showing the complete (left) and chimeric (right) absence of GFP and presence of mTagBFP2 signal specific to root cap cells. GFP is shown in green and mTagBFP2 in magenta. In left panels, white arrowheads indicate live root cap cells with nuclear mTagBFP2 signal covering the elongation zone, a typical phenotype observed when smb mutant seedlings are grown in presence of 1μM brassinazole. In right panels, red arrowheads indicate root cells still expressing GFP. Scale bars, 100 μm. **i**, Venn diagrams showing the number of plants displaying strong mTagBFP2 signal, *smb* and *gfp* phenotype out of 86 *pSMB(I)* (top) and 88 *pSMB(II)* (bottom) T1 seedlings. **j** Genotype analysis of pSMB (I) and pSMB (II) T2 lines by amplicon sequencing. The Heat map shows the percentage of indels for each target site. The phenotype observed for the corresponding seedling is indicated on the right panel. **k**, Frequency of the main mutations types in both pSMB (I) and pSMB (II) plants, for the six target sites. I1 to I3: 1- to 3-bp insertion, D1 to D6: 1- to 6-bp deletion, Others: bigger deletions (> 6-bp), insertions (> 3-bp) or more complex repair outcomes containing both insertions and deletions.

Forty-nine out of 96 *pPcUBI(I)*, and 52 out of 95 *pPcUBI(II)* T1 seedlings (selected by mRuby3 expression via a modified FAST system (Shimada *et al*., 2010; Decaestecker *et al*., 2019)) displayed both *gfp* and *smb* mutant phenotypes in roots, indicating simultaneous mutations (Figure 1c, d). Additionally, 44 out of 96 *pPcUBI(I)* and 45 out of 95 *pPcUBI(II)* T1 seedlings also lacked trichomes on the first two true leaves, revealing a high mutation frequency for *GL1*. Altogether, 79% of the *pPcUBI(I)* and 68% of the *pPcUBI(II)* T1 seedlings with at least one detectable knockout phenotype also showed triple *gfp smb gl1* mutant phenotypes. When selecting plants based on the loss of GFP, 90% of the *pPcUBI(I)* and 85% of the *pPcUBI(II)* T1 seedlings displayed triple mutant phenotypes, indicating a strong correlation of mutagenesis efficiencies in these three loci.

We quantified the indel frequency for the six targeted genes in 48 *pPcUBI(I)*, 47 *pPcUBI(II)* and a control NLS-GFP plant. The targeted loci were PCR amplified from root tips and amplicons were sequenced using Illumina sequencing (NGS) to determine the frequency and sequence of mutant alleles. Plants showing total or partial *gfp* and *smb* mutant phenotypes had high indel frequencies in *GFP* (27-100%) and *SMB* (38-98%) amplicons, as well as in all other target genes. Hierarchical clustering showed that transgenic T1 plants fell in two major classes that had either a high or a low level of mutagenesis for all six target genes (Figure 1e). In agreement with previous reports (Feng *et al*., 2019), 1-bp indels were the predominant repair outcome (50% to 80% and 1% to 15% respectively) and in-frame indels were rare (2% to 8%) (Figure 1f). Between 6% and 26% of mutations were bigger deletions (>6-bp), insertions (>3-bp) or more complex repair outcomes containing both insertions and deletions.

To test whether gRNA position within the vector has an effect, we compared indel frequencies for each target between the two constructs (Figure 1g). The overall percentage of indels between the *pPcUBI (I)* and *pPcUBI (II)* vectors was generally higher for *pPcUBI (II)*, though the difference was only significant for *GFP*. As the *GFP* gRNA was on the 5’ position in *pPcUBI (I)* and the 3’ position in *pPcUBI (II)*, one could hypothesize that the first gRNA position confers a higher indel efficiency than the last one. However, the percentage of indels for *SMB* with *pPcUBI (II)* (3’ position) was also higher than with *pPcUBI (I)* (5’ position), invalidating this hypothesis. As all other gRNAs had no substantial changes in indel frequencies, our data does not support a position effect in gRNA arrays, thus reducing the complexity of future experimental design.

We then tested whether six genes can be efficiently mutated in a tissue-specific context by making two vectors expressing Cas9-P2A-mTagBFP2 under the root cap-specific *pSMB* promoter with the same six gRNAs arranged in inverted orders (hereafter, *pSMB(I)* and *pSMB(II)). gfp* and *smb* mutant phenotypes were investigated using confocal microscopy in the T1 generation. Plants were grown in the presence of 1 μM brassinazole (BRZ) to facilitate *smb* phenotyping. This treatment leads to a root covered by living root cap cells in *smb* mutants (Fendrych *et al*., 2014) and was easily recognizable due to the presence of nuclear mTagBFP2 signal in living root cap cells (Figure 1h).

Thirty-two of 86 *pSMB(I)* and 46 of 88 *pSMB(II)* T1 seedlings showed both *gfp* and *smb* mutant phenotypes, as well as a strong mTagBFP2 signal specifically in root cap nuclei as determined by confocal microscopy (Figure 1i). In agreement with our previous report (Decaestecker *et al*., 2019), mTagBFP2 signal intensity could be used as a proxy for Cas9 expression and therefore the penetrance of *gfp* and *smb* knockout phenotypes. To determine mutagenesis efficiency in all target genes specifically in Cas9-expressing root cap cells, we collected root-tip protoplasts expressing mTagBFP2 (BFP^+^, Cas9 expressing cells) using fluorescence-activated cell sorting from T2 seedlings of ten *pSMB(I)* and eight *pSMB(II)* independent lines. We chose four *pSMB(II)* lines (19, 25, 35 and 48) with weak or chimeric *gfp* and *smb* T1 mutant phenotypes, and four *pSMB(I) and(II)* lines with highly penetrant *smb* and *gfp* T1 mutant phenotypes (Supplemental Figure 1).

The targeted loci were PCR amplified directly from sorted protoplast populations and analyzed by NGS. In *pSMB(I)* and *(II)* T2 seedlings coming from a T1 parent with strong *smb* and *gfp* phenotypes, the Cas9-expressing BFP^+^ populations had indel frequencies between 51% and 92% for all six target loci (Figure 1j). In contrast, the BFP^-^ GFP^+^ populations of these lines had indel frequencies of 1 to 26% (Supplemental figure 2). As expected, the BFP^+^ populations of the *pSMB(II)* lines that with weak or chimeric *gfp* and *smb* phenotypes in T1 had lower indel frequencies (2-50%), though they were still substantially higher than the frequencies in the BFP^-^ GFP^+^ populations (between 0% and 12%). Moreover, the relatively high indel frequencies in the BFP^-^ cells when compared with the same population in NLS-GFP seedlings are likely due to technical difficulties of sorting mTagBFP2 and GFP cells, and that some cells scored as mTagBFP2 negative are in fact positive (Supplemental figure 1). These results confirmed that in lines with high *GFP* and *SMB* mutagenesis activity, all six genes were simultaneously mutated with high efficiency.

Similarly to the ubiquitous lines, the alleles generated were largely consistent across events, with 1-bp indels being the predominant repair outcome (50-87% and 2-10%), in-frame insertion or deletions were rare (0-5%) and 3-21% of mutations were bigger indels (>3-bp and >6-bp) or combination of indels (Figure 1k).

In conclusion, we show that ubiquitous CRISPR and CRISPR-TSKO approaches allow fast and simultaneous disruption of six genes in the first transgenic generation with high efficiency. As mutation efficiencies over all six loci are correlated, we suggest the use of a target gene with an easy-to-score, nondetrimental loss-of-function phenotype as a proxy for highly mutagenized lines. As an alternative to endogenous genes (Li *et al*., 2020), loss of GFP in a reporter line can also be used as one such proxy. We foresee this approach to be a powerful tool to dissect genetic networks in model and crops species alike.

## Methods

### Vector construction

Cloning reactions were transformed into the *Escherichia coli* strain DH5α via heat-shock or One Shot ccdB Survival 2 T1R *E. Coli* Competent Cells (Thermo Fisher Scientific). Depending on the plasmid selectable marker, the cells were plated on LB (Lysogeny Broth) medium containing 100 mg.L^−1^ carbenicillin, 100 mg.L^−1^ spectinomycin or 10 mg.L^−1^ chloramphenicol. Colonies were verified via restriction enzyme digestion, and Sanger sequencing by Eurofins Scientific using the Mix2Seq service. All PCRs for cloning were performed with Q5 High-Fidelity DNA Polymerase (New England Biolabs) according to manufacturer’s instructions. Column and gel purifications were performed with Zymo-Spin II columns (Zymo Research) according to manufacturer instructions. Primers used for Cloning, Vector validation and genotyping are listed in supplemental table 1.

To create the *pPcUBI* (pFASTR-*pPcUBI*-Cas9-mTagBFP2-G7T-A-BsaI-ccdB-BsaI-G) vector, the entry modules pGG-A-PcUBIP-B, pGG-B-Linker-C, pGGC-Cas9-no-stop-D, pGG-D-mTagBFP2-E, pGG-E-G7T-F, and pGG-F-AarI-G were assembled into a pFASTRK-AG via a Golden Gate reaction as previously described (Decaestecker *et al*., 2019). After digestion and Sanger sequencing, an A-ccdB/CmR-G PCR fragment was inserted into the resulting vector via restriction enzyme digestion and ligation using AarI (Thermo Scientific™) and T4 DNA ligase (New England Biolabs) respectively as previously described (Decaestecker *et al*., 2019)

To clone the six gRNAs in a destination vector, for each target, a primer flanked by Gibson overhangs was cloned into a different BbsI digested unarmed entry vector by Gibson assembly. The six resulting armed gRNA modules were combined in the *pPcUBI* (pFASTR-*pPcUBI*-Cas9-mTagBFP2-G7T-A-BsaI-ccdB-BsaI-G) or *pSMB* (pFASTR-SMBP-Cas9-P2A-mTagBFP2-G7T-A-BsaI-ccdB-CmR-BsaI-G) destination vector via a Golden Gate reaction as previously described^3^. All plasmids reported here are listed in supplemental table 2 and available via the VIB-UGentCenter for Plant Systems Biology Gateway Vector website (https://gatewayvectors.vib.be/).

### Plant lines

The NLS-GFP line *(pHTR5:NLS-GFP-GUS)* was previously reported (Ingouff *et al*., 2017) (NASC collection N2109788).

### Plant Transformation and Selection

Plant vectors were transformed in *Agrobacterium tumefaciens* GV3101 pMP90 strain by electroporation. *pHTR5:NLS-GFP-GUS* Arabidopsis line was transformed via the floral-dip method^15^. FASTR-positive T1 transgenic seeds were selected under a Leica M165FC fluorescence stereomicroscope using the DSR fluorescence filter (excitation: 510–560 nm; emission 590–650 nm).

Selected seeds were surface sterilized with 70% ethanol for 1 min and 3% sodium hypochlorite /0,01% Tween-20 for 10 min and washed 5 times with sterile water. Seeds were plated on 2.15 g L^−1^ MS salts (Duchefa M0221) with 0.1 g L^−1^ MES.H2O medium containing 8g L^−1^ plant agar and stratified for 2 days at 4°C. Seedlings were grown in vertically oriented square petri plates inside 21°C growth chambers under continuous light from SpectraluxPlus NL36W/840 Plus (Radium Lampenwerk) fluorescent bulbs.

After confocal observation, T1 plants were transferred to Jiffy-7 pellets and grown in a greenhouse at 21°C under a 16-h day regime (100 W m^−2^ photosynthetically active radiation) from natural light complemented with 600-W GreenPower (Philips) high-pressure sodium lightbulbs.

### Genotyping

For the *pPcUBI* lines, the root tip of each T0 seedling at 7DAG was used to perform PCR directly, without DNA purification, using Thermo Scientific™ Phire™ Plant Direct PCR Master Mix kit and following the manufacturer’s protocol. Briefly, each root tip was transferred into a well of 96-well plate containing 40 μL of dilution buffer and roughly crushed using a 200μL pipette tip. For the TSKO lines the mTagBFP2-positive protoplasts were directly sorted into Phire™ Plant Direct PCR Master Mix dilution buffer.

For each sample, the six genomic regions flanking the target site of each gene were amplified with specific primers (supplemental table 1) and the PCR products were analyzed via agarose gel electrophoresis.

For the *pPcUBI*, 144 PCR products, resulting from the amplification of the 6 targets in 24 plants, were pooled. For the TSKO, 48 PCR products, resulting from the amplification of the 6 targets in 8 protoplast-sorted samples, were pooled. The PCR pool cleanup was done by column purification with a DNA Clean and Concentrator kit (Zymo Research). The purified samples were sent for EZ-amplicon sequencing using Illumina-based technology (2×250 bp reads) by Genewiz.

### Mutagenesis analysis

Amplicon sequencing analysis was performed within Galaxy (https://usegalaxy.eu/). Paired-end reads were demultiplexed using Je-Demultiplex^16^, merged with FLASH^17^, and trimmed and filtered for quality using Trimmomatic^18^ using default parameters. Merged reads retained after filtering were used to identify and quantify the percentage of mutation at each target site using AGESeq with a mismatch rate cut-off of 0.1 and without any abundance cut-off^19^.

### Confocal microscopy

Root tips of 6DAG T1 seedlings grown under continuous light on 2.15 g L^-^1 MS salts with 0.1 g L^−1^ MES.H2O with (for *pSMB (I)* and *(II))* or without (for *pPcUBI (I)* and *(II))* 1 μM Brassinazole medium were imaged using an Ultra View Vox Spinning disc confocal imaging system (PerkinElmer) mounted on an Eclipse Ti inverted microscope (Nikon) using a 512×512 Hamamatsu ImagEM C9100-13 EMccd cameras to image.

Samples were stained with 10 mg mL^−1^ propidium iodide (PI) in 0.215 gL^−1^ MS salts (Duchefa M0221) with 0.01 g L^−1^ MES.H2O medium. GFP was excited at 488 nm and acquired with an emission window between 500 and 530 nm. PI was imaged with 561 nm excitation light and an emission window between 570nm and 625nm. mTagBFP2 was imaged with 405 nm excitation light and an emission window between 454nm and 496nm.

### Protoplast Preparation and Cell Sorting

Arabidopsis root protoplasts were prepared as previously described^20^. In brief, for *pSMB (I)* and *(II)* lines, root tips of around 600 6DAG seedlings grown under continuous light on 0.43 g L^−1^ MS salts with 94 μM MES.H2O medium containing 1 μM Brassinazole were incubated in protoplasting solution consisting of 1.25% (w/v) cellulase (Yakult), 0.3% (w/v) macerozyme (Yakult), 0.4 M mannitol, 20 mM MES, 20 mM KCl, 0.1% (w/v) BSA, and 10 mM CaCl2 at pH 5.7 for 3 h. The protoplast solutions were filtered through a 40μm cell strainer into a 15mL round-bottom tube and centrifuged at 150g for 10 min. The supernatant was discarded and protoplasts were recovered in 4°C resuspension buffer (buffer with the same composition as protoplasting buffer but without cellulase and macerozyme).

Root tip protoplasts were sorted into 1.5-mL Eppendorf tubes containing 25μL of Thermo Scientific™ Phire™ Plant Direct PCR Master Mix dilution buffer using a BD FACSMelody equipped with three lasers (405 nm, 488 nm, and 561 nm).

### Accession numbers

Gene models used in this article can be found in the Arabidopsis Genome Initiative database under the following accession numbers: *SMB* (AT1G79580); *EXI1* (AT2G14095); *GL1* (AT3G27920); *ARF7* (AT5G20730); *ARF19* (AT1G19220).

Seeds for the line *pHTR5:NLS-GFP-GUS* are available from the Nottingham Arabidopsis Stock Centre (NASC), ID N2109788.

## Acknowledgements

This work was supported by FWO (G041118N) and ERC (639234 and 864952). We thank Dr. Jonas Blomme for providing the *pPcUBI* and pGG-D-mTagBFP2-E vectors. We thank Gert Van Isterdael and Julie Van Duyse (VIB Flow Core) for support and access to the instrument park. We kindly thank Dr. Zongcheng Lin, Dr. Nicolas Doll, Ward Decaestecker, Dr. Christophe Gaillochet, Ania Lukasiewicz, Dr. Jonas Blomme and Ward Develtere for their critical review of the manuscript.

## Author contributions

N.B., R.A.B., M.K.N. and T.B.J. designed the experiments and wrote the manuscript. N.B. and R.A.B. performed the experiments and analyzed the data.

## References

Babicki S, Arndt D, Marcu A, Liang Y, Grant JR, Maciejewski A and Wishart DS, (2016). Heatmapper: web-enabled heat mapping for all. Nucleic acids research, 44:147–153.

Bargmann, BOR & Birnbaum KD (2010) Fluorescence activated cell sorting of plant protoplasts. J. Vis. Exp., 36:e1673.

Bolger AM, Lohse M and Usadel B (2014) Trimmomatic: A flexible trimmer for Illumina sequence data. Bioinformatics, 30:2114–2120.

Clough SJ and Bent AF (1998) Floral dip: A simplified method for Agrobacterium-mediated transformation of Arabidopsis thaliana. Plant J., 16:735–743 (1998).

Decaestecker W, Buono RA, Pfeiffer ML, Vangheluwe N, Jourquin J, Karimi M, Van Isterdael G, Beeckman T, Nowack MK and Jacobs TB, (2019). CRISPR-TSKO: a technique for efficient mutagenesis in specific cell types, tissues, or organs in Arabidopsis. The Plant Cell, 31:2868–2887.

Fendrych M, Van Hautegem T, Van Durme M, Olvera-Carrillo Y, Huysmans M, Karimi M, Lippens S, Guérin CJ, Krebs M, Schumacher K and Nowack MK, (2014). Programmed cell death controlled by ANAC033/SOMBRERO determines root cap organ size in Arabidopsis. Current Biology, 24:931–940.

Feng Z, Mao Y, Xu N, Zhang B, Wei P, Yang DL, Wang Z, Zhang Z, Zheng R, Yang L and Zeng L, (2014). Multigeneration analysis reveals the inheritance, specificity, and patterns of CRISPR/Cas-induced gene modifications in Arabidopsis. Proceedings of the National Academy of Sciences, 111:4632–4637.

Girardot C, Scholtalbers J, Sauer S, Su SY and Furlong EEM (2016) Je, a versatile suite to handle multiplexed NGS libraries with unique molecular identifiers. BMC Bioinformatic,s 17:4–9.

Herman PL and Marks MD, (1989). Trichome development in Arabidopsis thaliana. II. Isolation and complementation of the GLABROUS1 gene. The Plant Cell, 1:1051–1055.

Ingouff, M. et al. (2017) Live-cell analysis of DNA methylation during sexual reproduction in arabidopsis reveals context and sex-specific dynamics controlled by noncanonical RdDM. Genes Dev. 31:72–83.

Kurata M, Wolf NK, Lahr WS, Weg MT, Kluesner MG, Lee S, Hui K, Shiraiwa M, Webber BR and Moriarity BS, (2018). Highly multiplexed genome engineering using CRISPR/Cas9 gRNA arrays. PLoS one, 13:0198714.

Li R, Vavrik C and Danna CH, 2020. Proxies of CRISPR/Cas9 Activity To Aid in the Identification of Mutagenized Arabidopsis Plants. G3: Genes, Genomes, Genetics, 10:2033–2042.

Liang Y, Eudes A, Yogiswara S, Jing B, Benites VT, Yamanaka R, Cheng-Yue C, Baidoo EE, Mortimer JC, Scheller HV and Loqué D, (2019). A screening method to identify efficient sgRNAs in Arabidopsis, used in conjunction with cell-specific lignin reduction. Biotechnology for biofuels, 12:1–15.

Magoč T and Salzberg SL (2011) FLASH: Fast length adjustment of short reads to improve genome assemblies. Bioinformatics 27:2957–2963.

Mao Y, Botella J R, Liu Y and Zhu JK Gene editing in plants: Progress and challenges (2019). Natl. Sci. Rev. 6:421–437.

Shimada TL, Shimada T and Hara-Nishimura I (2010) A rapid and non-destructive screenable marker, FAST, for identifying transformed seeds of Arabidopsis thaliana. Plant J. 61:519–528

Wang X, Ye L, Lyu M, Ursache R, Löytynoja A and Mähönen AP, 2020. An inducible genome editing system for plants. Nature plants, 6:766–772.

Xue LJ and Tsai CJ (2015) AGEseq: Analysis of genome editing by sequencing. Mol. Plant 8:1428–1430.

Zhang Y, Malzahn AA, Sretenovic S. and Qi Y, (2019). The emerging and uncultivated potential of CRISPR technology in plant science. Nature Plants, 5:778–794.

Zhang Z, Mao Y, Ha S, Liu W, Botella JR and Zhu JK, (2016). A multiplex CRISPR/Cas9 platform for fast and efficient editing of multiple genes in Arabidopsis. Plant cell reports, 35:1519–1533.

