## Supplemental_data_1 for "Efficient simultaneous mutagenesis of multiple genes in specific plant tissues by multiplex CRISPR"

*pSMB (I)* line 50

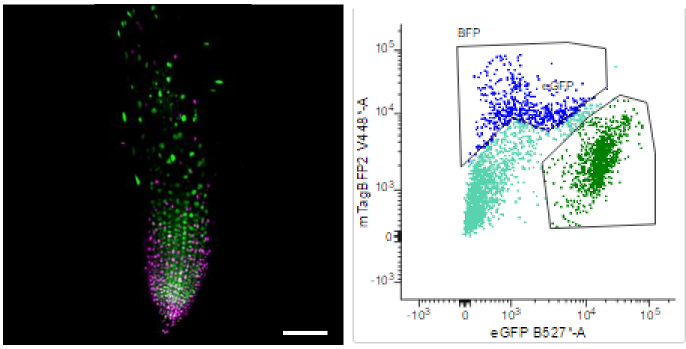


*pSMB (I)* line 24

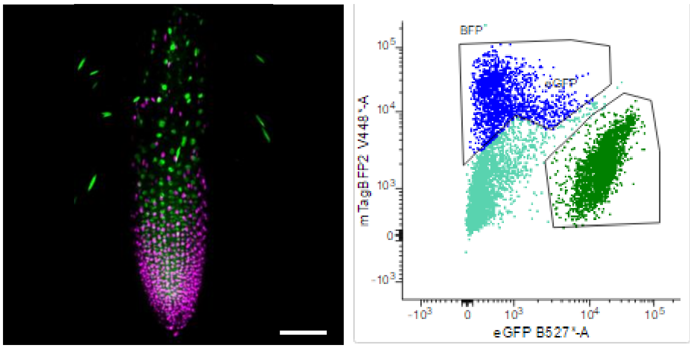


*pSMB (I)* line 75

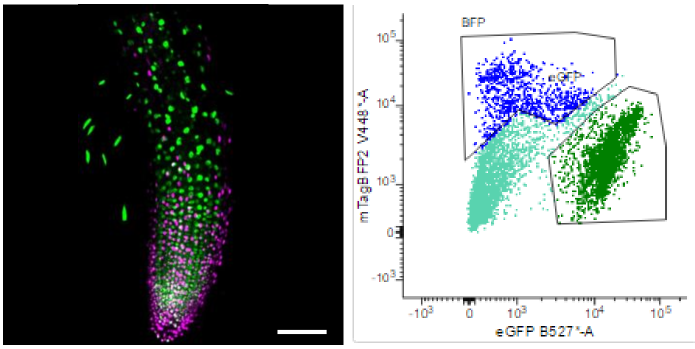


*pSMB (I)* line 41

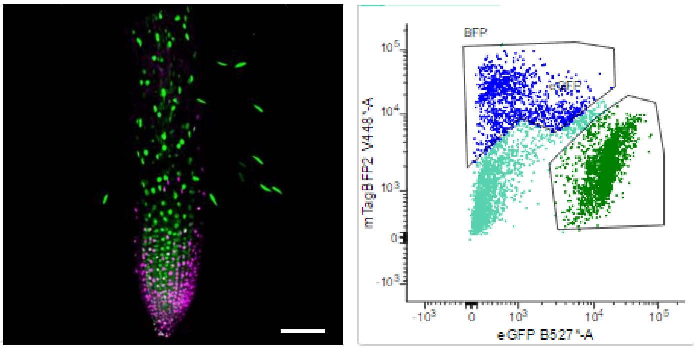


*pSMB (II)* line 21

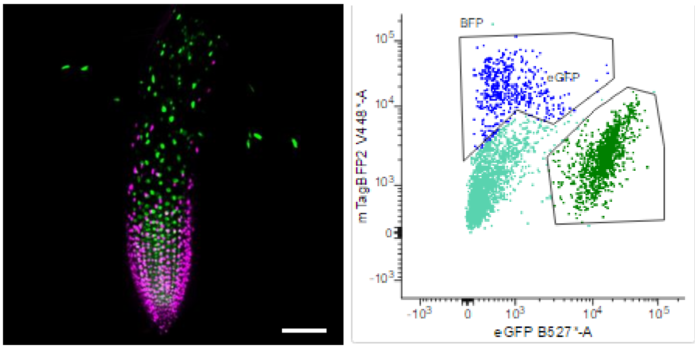


*pSMB (II)* line 8

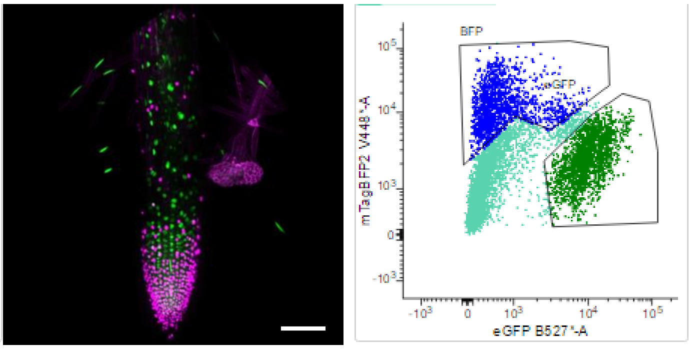


*pSMB (II)* line 35

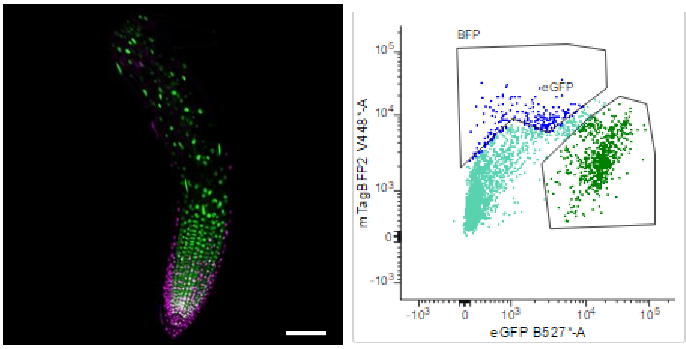


*pSMB (II)* line 19

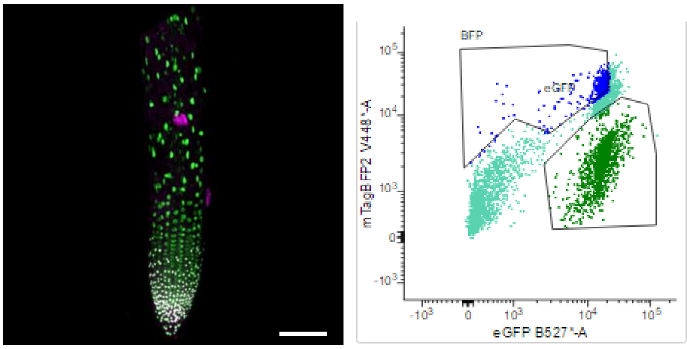


**Supplemental figure 1. Fluorescence-activated Cell Sorting of protoplasts used in genotyping.**
Left panels are maximum intensity projections of *pSMB (I)* and *pSMB (II)* T1 mother lines. GFP is shown in green and Cas9-P2A-mTagBFP2 in magenta.
Right panel are the 50.000 events recorded during protoplast sorting showing the gating strategies. $BFP^+$ and $BFP^-$ $GFP^+$ collected populations were used for DNA extraction and molecular analysis.

*pSMB (I)* line 3

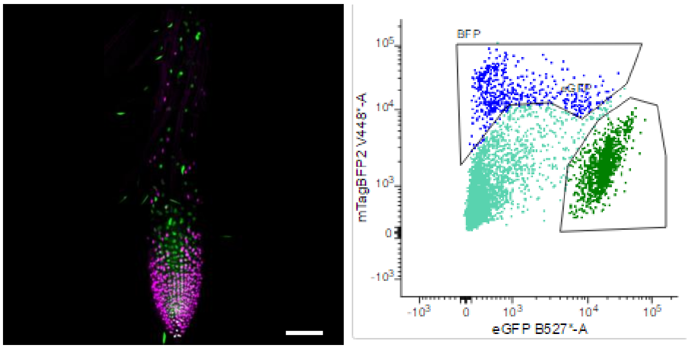


*pSMB (I)* line 52

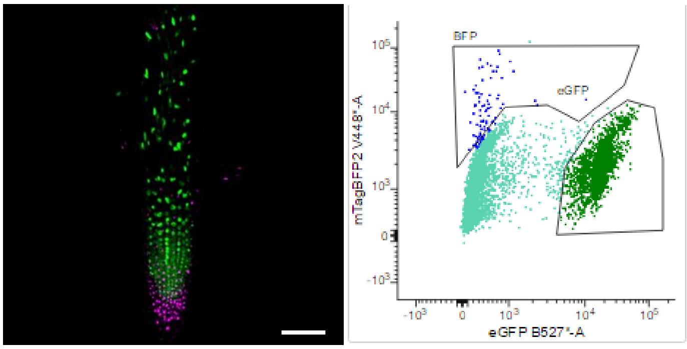


*pSMB (I)* line 45

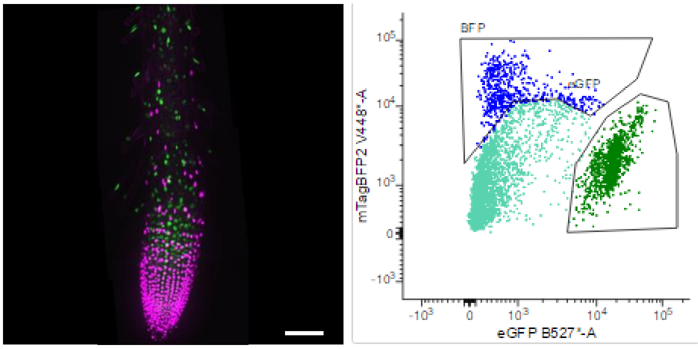


*pSMB (I)* line 58

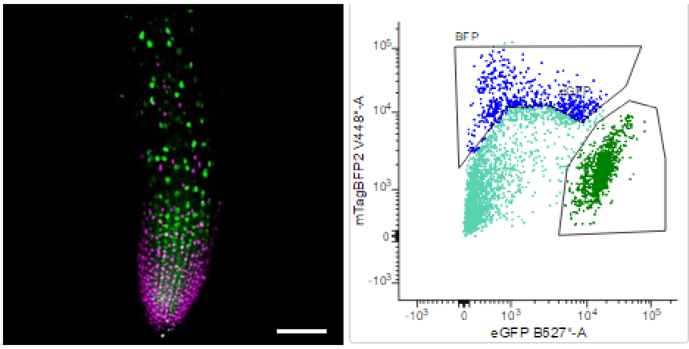


*pSMB (I)* line 48

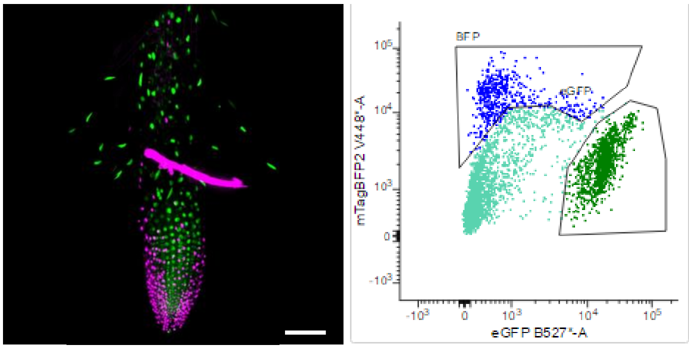


*pSMB (I)* line 59

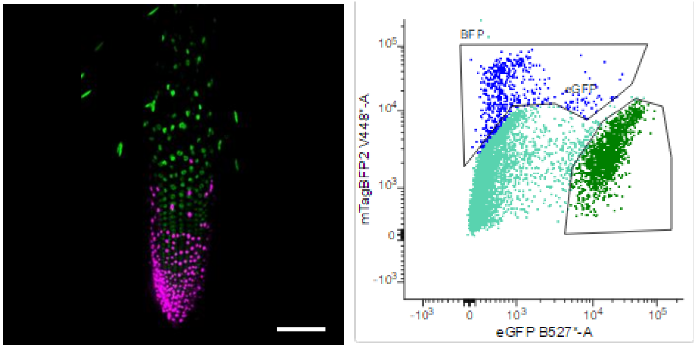


*pSMB (II)* line 25

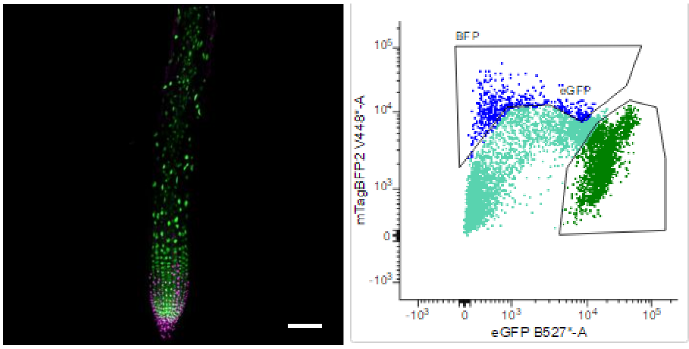


*pSMB (II)* line 48

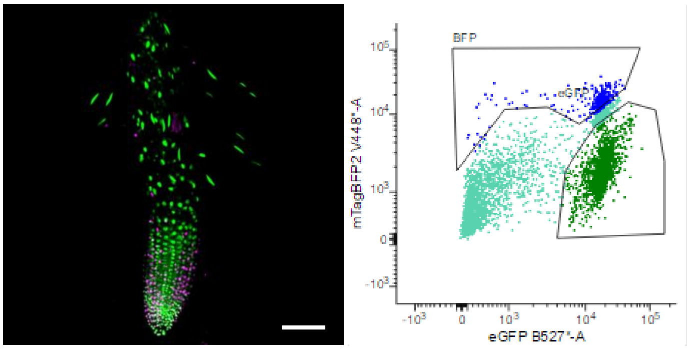


**Supplemental figure 1. Continued**

*pSMB (II)* line 30

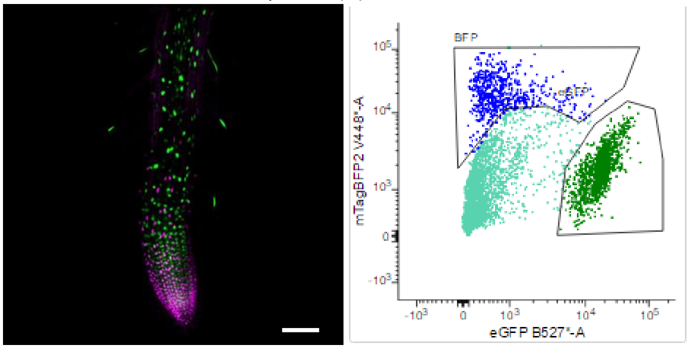


*pSMB (II)* line 55

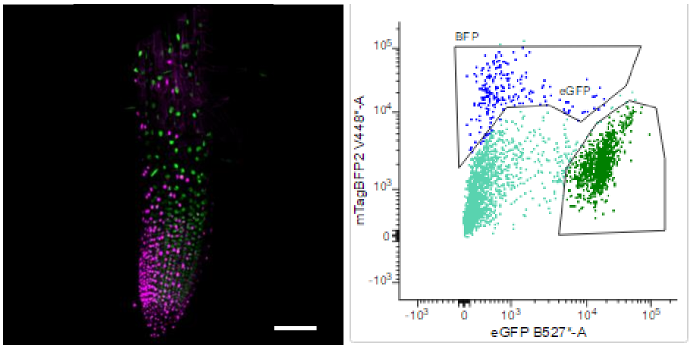


*pH3.3::NLS-GFP*

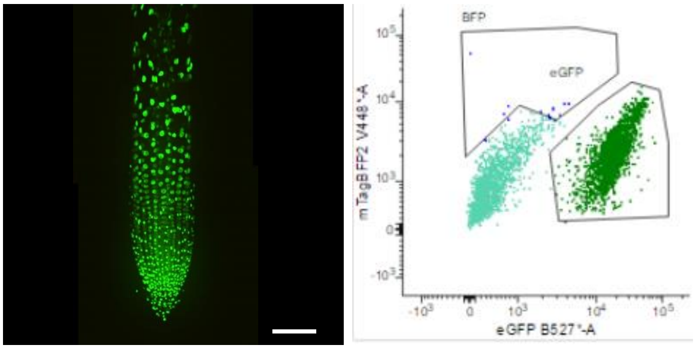


Col-0

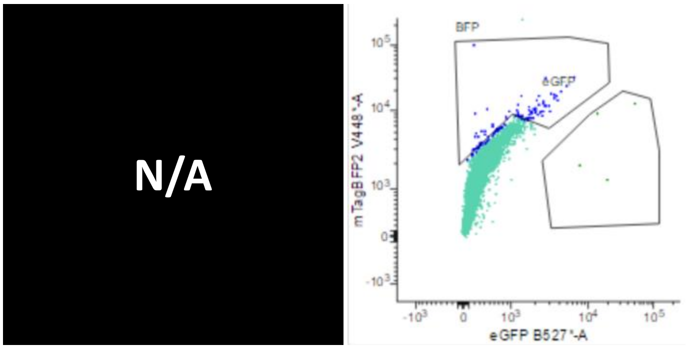


**Supplemental figure 1. Continued**

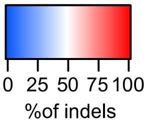


| Percentage of indels | | | | | | | | Smb$^{-}$ | GFP signal | |
|---|---|---|---|---|---|---|---|---|---|---|
| SMB | EXI1 | GL1 | ARF7 | ARF19 | GFP | Line | Sorting Gate | Yes | No | Chimeric |
| 74 | 63 | 80 | 72 | 83 | 82 | 48 | BFP$^{+}$ | + | + | |
| 1 | 2 | 5 | 4 | 7 | 8 | | BFP$^{-}$ GFP$^{+}$ | | | |
| 70 | 71 | 72 | 72 | 70 | 81 | 24 | BFP+ | + | + | |
| 11 | 3 | 10 | 7 | 30 | 3 | | BFP- GFP+ | | | |
| 79 | 56 | 82 | 73 | 61 | 71 | 3 | BFP+ | + | + | |
| 4 | 4 | 2 | 3 | 8 | 7 | | BFP- GFP+ | | | |
| 82 | 65 | 82 | 61 | 60 | 70 | 41 | BFP+ | + | + | |
| 14 | 11 | 13 | 13 | 18 | 13 | | BFP- GFP+ | | | |
| 74 | 62 | 65 | 85 | 63 | 69 | 59 | BFP+ | + | + | |
| 2 | 2 | 6 | 2 | 0 | 2 | | BFP- GFP+ | | | |
| 54 | 72 | 87 | 84 | 59 | 67 | 75 | BFP+ | + | + | |
| 16 | 16 | 19 | 12 | 10 | 3 | | BFP- GFP+ | | | |
| 69 | 72 | 88 | 58 | 68 | 61 | 45 | BFP+ | + | + | |
| 31 | 23 | 12 | 21 | 26 | 19 | | BFP- GFP+ | | | |
| 69 | 75 | 69 | 58 | 82 | 59 | 52 | BFP+ | + | + | |
| 5 | 1 | 2 | 3 | 3 | 1 | | BFP- GFP+ | | | |
| 52 | 61 | 68 | 51 | 72 | 65 | 58 | BFP+ | + | + | |
| 5 | 3 | 7 | 4 | 4 | 7 | | BFP- GFP+ | | | |
| 58 | 67 | 52 | 63 | 51 | 61 | 50 | BFP+ | + | + | |
| 17 | 24 | 16 | 15 | 13 | 16 | | BFP- GFP+ | | | |
| 73 | 73 | 81 | 77 | 84 | 92 | 21 | BFP+ | + | + | |
| 19 | 13 | 8 | 18 | 10 | 12 | | BFP- GFP+ | | | |
| 84 | 77 | 91 | 84 | 77 | 84 | 30 | BFP+ | + | + | |
| 3 | 3 | 1 | 3 | 6 | 5 | | BFP- GFP+ | | | |
| 62 | 66 | 86 | 56 | 83 | 59 | 8 | BFP+ | + | + | |
| 17 | 13 | 13 | 5 | 1 | 12 | | BFP- GFP+ | | | |
| 68 | 54 | 72 | 55 | 40 | 72 | 55 | BFP+ | + | + | |
| 3 | 3 | 2 | 1 | 6 | 1 | | BFP- GFP+ | | | |
| 42 | 42 | 36 | 36 | 31 | 50 | 25 | BFP+ | | | + |
| 10 | 10 | 10 | 5 | 9 | 7 | | BFP- GFP+ | | | |
| 36 | 43 | 42 | 43 | 34 | 28 | 35 | BFP+ | | | + |
| 5 | 4 | 3 | 9 | 0 | 9 | | BFP- GFP+ | | | |
| 17 | 14 | 18 | 12 | 19 | 15 | 19 | BFP+ | | | + |
| 6 | 1 | 3 | 0 | 0 | 3 | | BFP- GFP+ | | | |
| 46 | 17 | 11 | 11 | 2 | 17 | 48 | BFP+ | + | | + |
| 7 | 6 | 9 | 8 | 8 | 12 | | BFP- GFP+ | | | |
| 2 | 2 | 1 | 0 | 0 | 0 | GFP (1) | BFP- GFP+ | N/A | N/A | N/A |
| 1 | 0 | 0 | 1 | 1 | 1 | GFP (2) | BFP- GFP+ | N/A | N/A | N/A |
| 0 | 0 | 0 | 0 | 1 | 0 | GFP (3) | BFP- GFP+ | N/A | N/A | N/A |
| 1 | 1 | 2 | 2 | 1 | 1 | GFP (4) | BFP- GFP+ | N/A | N/A | N/A |

**Supplemental figure 2. Genotype analysis of *pSMB (I)* and *pSMB (II)* T2 lines by amplicon sequencing.**
The Heat map is showing the percentage of indels for each target site in each sorted population (BFP$^{+}$ and BFP$^{-}$ GFP$^{+}$) for each line.
The phenotype observed for the corresponding T1 motherline is indicated on the right panel.
